# Myofiber injury induces capillary disruption and regeneration of disorganized microvascular networks

**DOI:** 10.1101/2021.08.02.454805

**Authors:** Nicole L. Jacobsen, Charles E. Norton, Rebecca L. Shaw, DDW Cornelison, Steven S. Segal

**Affiliations:** Medical Pharmacology and Physiology, University of Missouri, Columbia, MO 65212; Biological Sciences, University of Missouri, Columbia, MO 65212; Christopher S. Bond Life Sciences Center, University of Missouri, Columbia, MO 65212; Dalton Cardiovascular Research Center, University of Missouri, Columbia, MO 65212

**Author notes:** Correspondence: Steven S. Segal, PhD, University of Missouri, Medical Pharmacology and Physiology, MA415 Medical Sciences Building, 1 Hospital Drive, Columbia, MO 65212.

**Keywords:** capillaries, microcirculation, myofiber, skeletal muscle regeneration

## Abstract

Myofibers regenerate following injury, however the microvasculature must also recover to restore skeletal muscle function. We aimed to define the nature of microvascular damage and repair during skeletal muscle injury and regeneration induced by BaCl_2_. To test the hypothesis that microvascular disruption occurred secondary to myofiber injury in mice, isolated microvessels were exposed to BaCl_2_ or the myotoxin was injected into the gluteus maximus (GM) muscle. In isolated microvessels, BaCl_2_ depolarized smooth muscle cells and endothelial cells while increasing [Ca^2+^]_i,_ but did not elicit cell death. At 1 day post injury (dpi) of the GM, capillary fragmentation coincided with myofiber degeneration while arteriolar and venular networks remained intact; neutrophil depletion before injury did not prevent capillary damage. Perfused capillary networks reformed by 5 dpi in association with more terminal arterioles and were dilated through 10 dpi; with no change in microvascular area or branch point number in regenerating networks, fewer capillaries aligned with myofibers and capillary networks were no longer organized into microvascular units. By 21 dpi, capillary orientation and organization had nearly recovered to that in uninjured GM. We conclude that following their disruption secondary to myofiber damage, capillaries regenerate as disorganized networks that remodel while regenerated myofibers mature.

## Introduction

Skeletal muscle regeneration is an intricate process that requires the activation, proliferation, and differentiation of resident stem cells called satellite cells (1). However, the restoration of intact, functional muscle requires the coordinated recovery of additional cell types and tissue components during myogenesis, particularly the microcirculation. When compared to the well-defined molecular and cellular events of myofiber degeneration and regeneration [for review, see (2)], little is known of how skeletal muscle injury and regeneration affect its microvascular supply.

The microvasculature of skeletal muscle consists of arterioles, capillaries, and venules comprising networks of branches arranged in series and in parallel (3). The microcirculation delivers oxygen and nutrients to myofibers while removing cellular debris and products of metabolism. To meet these physiological demands, the microcirculation responds acutely by regulating local blood flow [e.g., functional hyperemia in response to muscle contraction, (4)] and adapts to chronic use by modifying network morphology [e.g., increased capillarization, arteriogenesis; (5, 6)].

To study muscle injury and regeneration in mice, intramuscular injection of the myotoxic agent BaCl_2_ induces reproducible damage while sparing sufficient satellite cells to support myofiber regeneration (7). As shown in the gluteus maximus muscle (GM), BaCl_2_ injection also damages capillaries (8, 9), which undergo fragmentation within 1 day post injury (dpi), thereby eliminating local perfusion and solute transport. BaCl_2_-induced myofiber death occurs through depolarization of the sarcolemma leading to Ca^2+^ overload, membrane disruption, and proteolysis (8). However, it is unknown how BaCl_2_ results in capillary fragmentation *in vivo*. While freeze injury and injection of snake venom toxins also disrupt capillaries accompanied by with myofiber damage (7), it is unknown whether capillary fragmentation is a direct effect of the initial insult by BaCl_2_ or is a consequence of myofiber disruption. Nor has it been determined whether arteriolar and venular networks, which supply and drain capillary networks, are disrupted in the manner shown for capillaries.

Ischemic injury to skeletal muscle is followed by robust angiogenesis with an increase in capillary-to-myofiber ratio (7, 10). Following injury with BaCl_2_, endothelial sprouts appear within 2-3 dpi, with the ensuing regeneration of capillary networks restoring local perfusion by 5 dpi (8, 9). At this time, arteriolar networks are abnormally dilated, with recovery of blood flow control (vasomotor tone, dilation, and constriction) occurring by 21 dpi in the mouse GM as regenerating myofibers mature (8). Nevertheless, the cellular dynamics of revascularization and microvascular remodeling during myofiber regeneration are poorly understood. In this study we tested the hypotheses that 1) BaCl_2_ induces death of microvascular endothelial cells (ECs) and smooth muscle cells (SMCs) by triggering Ca^2+^ overload; and 2) capillaries proliferate during early regeneration with networks remodeling as regenerating myofibers mature.

## Results

Previous studies evaluating microvascular injury did not resolve whether damage and loss of perfusion were a direct effect of BaCl_2_ on vascular cells or was secondary to disruption of myofibers and leukocyte infiltration (7–9). Therefore, to evaluate the effect of BaCl_2_ on microvascular ECs and SMCs independent of surrounding myofibers or inflammation, superior epigastric arteries (SEAs) having a single layer of each cell type were isolated and exposed to BaCl_2_ *in vitro*. SMCs were studied in the wall of intact vessels and ECs were evaluated in endothelial tubes following dissociation of SMCs (11).

### BaCl_2_ induces depolarization and increases Ca^2+^ in SMCs causing vasoconstriction

Adding 1.2% BaCl_2_ during superfusion of SEAs progressively depolarized SMCs from −48 ± 5 mV (baseline) to −19 ± 2 mV over several min (Fig 1A & B). In endothelial tubes, BaCl_2_ depolarized ECs from −41 ± 1 mV (baseline) to −20 ± 2 mV over a similar time course (Fig 1B). Electrical responses plateaued in 7 ± 1 min.

**Figure 1.**
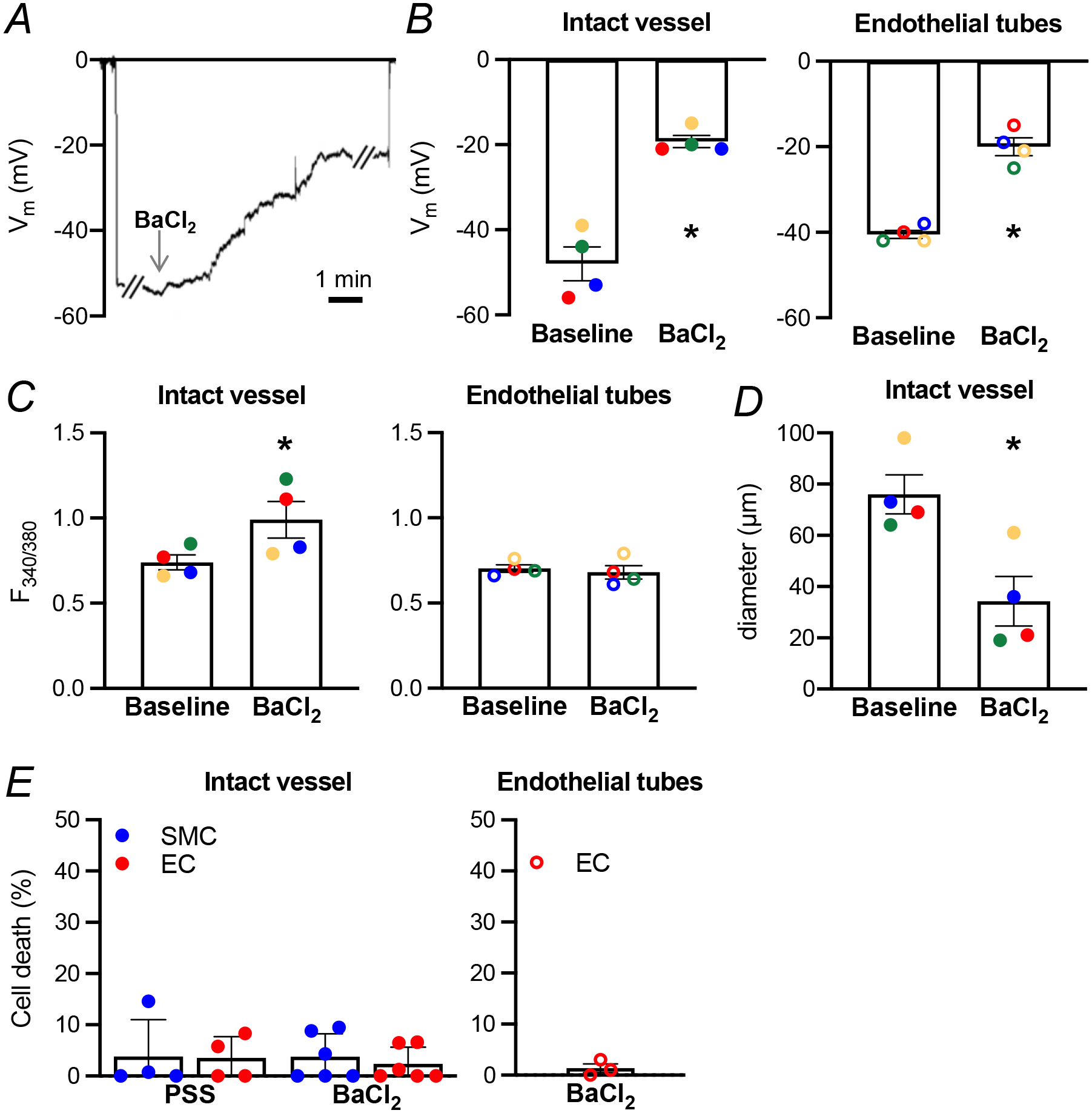
BaCl_2_ induces depolarization, calcium influx, and vasoconstriction in microvessels without cell death. **A)** Representative recording of V_m_ from a SMC in a pressurized SEA. Addition of BaCl_2_ initiated progressive depolarization that stabilized after several min. **B)** BaCl_2_ depolarizes SMCs in intact vessels (left) and ECs in endothelial tubes (right). For SMCs, ΔV_m_ = −29 mV ± 3 mV (P = 0.002, n = 4); for ECs, ΔV_m_ = −21 mV ± 2 mV (P = 0.001, n = 4). **C)** Cytosolic calcium concentration increases in SMCs of intact microvessels (P = 0.03, n = 4) but not in ECs of endothelial tubes (P = 0.31, n = 4) during exposure to BaCl_2_. Summary data of [Ca^2+^]_i_ measured with fura 2 fluorescence (F_340/380_) before and during BaCl_2_ application. **D)** Intact microvessels constrict 55% after 1 h exposure to BaCl_2_. Paired average diameter before (baseline) and after BaCl_2_ addition (P = 0.0007, n = 4). **E)** BaCl_2_ exposure does not induce cell death of SMCs or ECs *in vitro* (n = 4-6 vessels) Summary data presented as mean ± SE. Each color represents a separate paired experiment.

In SMCs, membrane depolarization is accompanied by a rise in [Ca^2+^]_i_ through activation of voltage gated (L-type) Ca^2+^ channels in the plasma membrane (12). Corresponding to depolarization (Fig 1B), exposure to BaCl_2_ increased SMC [Ca^2+^]_i_ (Fig 1C) and constricted SEAs from 76 ± 8 µm (baseline) to 34 ± 10 µm BaCl_2_ (Fig 1D). In the absence of voltage gated Ca^2+^ channels in ECs (13), depolarization with BaCl_2_ had no effect on EC [Ca^2+^]_i_. Controls performed without BaCl_2_ confirmed that V_m_ and [Ca^2+^]_i_ of SMCs in pressurized SEAs and of ECs in endothelial tubes remained stable for the duration of recordings (14–16).

### BaCl_2_ does not directly cause microvascular cell death

Exposure of the mouse extensor digitorum longus (EDL) muscle to 1.2% BaCl_2_ promptly depolarized myofibers and increased [Ca^2+^]_i_, leading to disruption of the sarcolemma and proteolysis culminating in cell death within 1 h (8). We questioned whether BaCl_2_ affected microvascular cells in a similar manner. Contrary to our hypothesis, 1 h exposure to 1.2% BaCl_2_ was not lethal to SMCs or ECs in intact vessels or to ECs in endothelial tubes (Fig 1E). Furthermore, extending BaCl_2_ exposure to 3 h had no further effect on EC or SMC viability (n = 3, not shown). These data suggest that, rather than a direct effect of BaCl_2_, the adverse microenvironment within an intact muscle created by myofiber injury and degeneration leads to disruption of capillary ECs.

### Arterioles and venules are spared from BaCl_2_-induced muscle injury

Capillaries are fragmented and perfusion is abolished within 24 h of muscle injury (7–9). At this early timepoint, leakage of a 70 kDa dextran from residual microvessels suggested a loss of structural integrity of arterioles or venules (9). To investigate whether pre- and post-capillary microvessels are also damaged by muscle injury, the GM of male C57Bl/6J mice was injured by injection of 75 µL of 1.2% BaCl_2_ under the muscle (9). Whole mount immunostaining of the GM with Myh11 labelled SMCs that encircle arterioles and venules embedded within skeletal muscle. While other cell types may express Myh11, they are not localized to the abluminal surface of larger caliber microvessels thereby enabling positive identification of SMCs (17, 18). As seen for isolated microvessels exposed to BaCl_2_ (Fig 1E), the structural integrity of SMCs within arteriolar networks of the GM remained essentially intact at 1 dpi, though infrequent damage was observed (Fig 2). Continuous SMC coverage and arteriolar segments with uniform diameter was not different at 1 dpi compared to uninjured muscle. The integrity of venular segments and their SMCs was also preserved (Fig 2A & B). Our quantitative analyses therefore centered on capillary and precapillary (resistance) networks.

**Figure 2.**
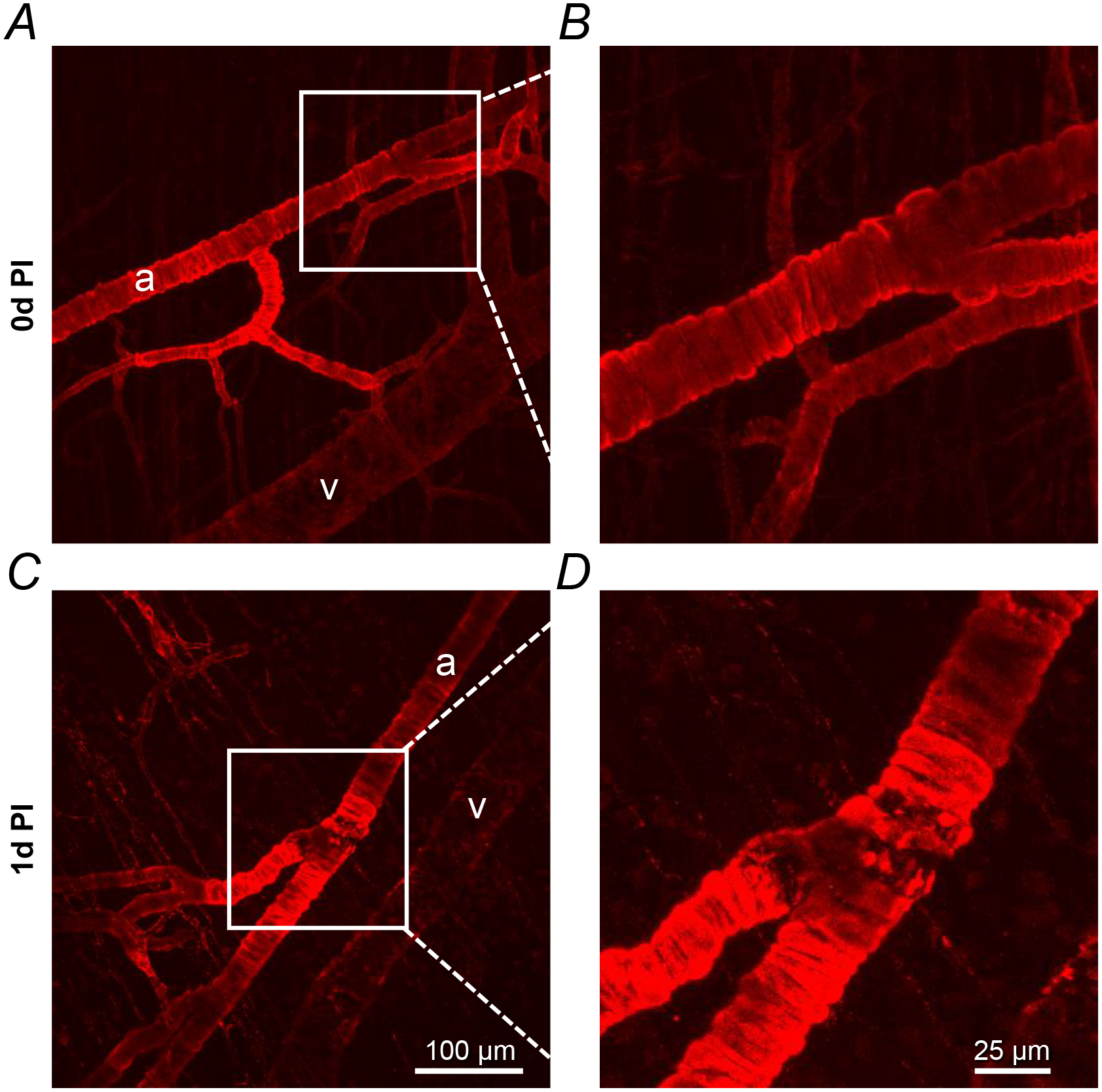
BaCl2 injection does not disrupt SMCs of arterioles (a) or venules (v) embedded in the GM. Representative maximum projection confocal z-stacks showing SMCs identified by Myh11 expression at **(A-B)** 0 dpi and **(C-D)** 1 dpi. Note example of rare local damage to SMCs at 1 dpi. Boxed regions within left panels are enlarged in right panels. Red = immunostaining for Myh11. Images are representative of n = 7 tissue regions from 3 mice.

### Role of neutrophils in BaCl_2_-induced capillary damage

Muscle injury initiates a stereotypical inflammatory response, with neutrophils invading within 1-2 h of injury and peaking at 12-24 h post injury (19). Neutrophils release cytolytic and cytotoxic molecules that may damage other resident cell types, along with cytokines that attract monocytes and macrophages to remove cellular debris (20). Given the disruption of capillaries following intramuscular injection of BaCl_2_ (7, 8) compared to the integrity of ECs exposed to BaCl_2_ *in vitro* (Fig 1E), we tested whether invading neutrophils are necessary for capillary fragmentation after BaCl_2_ injury. Differential blood counts determined that circulating neutrophils comprised 12.6 ± 1.8% (n = 4) of white blood cells in uninjured male C57BL/6J mice. At 1 dpi, neutrophils increased to 49.5 ± 3.2% of circulating white blood cells (P <0.0001). Intraperitoneal injections of anti-Ly6G ab prior to BaCl_2_ injury reduced circulating neutrophils at 0 dpi and at 1 dpi (4.3 ± 3.0% and 2.9 ± 2.0%, respectively; P <0.0001; n = 4 mice per group).

Using an EC-specific Cre driver [Cdh5-Cre^ERT2^; (21)] and tamoxifen-induced recombination of the Rosa26^mTmG^ locus to genetically label the endothelium with membrane-bound eGFP (Cdh5-mTmG mice), we evaluated capillary network structure in whole mount preparations of the GM. The orderly arrangement of capillary networks that course along myofibers in uninjured muscle (Fig 3A) was not impacted by neutrophil depletion alone. Following BaCl_2_ injury, capillaries were fragmented at 1 dpi (Fig 3A), which explains the loss of perfusion at this time (9). While neutrophil depletion did not prevent this capillary damage, there was a trend for preserving tissue area occupied by capillaries (Fig 3B, P = 0.28). As an index of capillary fragmentation with fewer pathways for blood flow, the maximum continuous length of networks was decreased at 1 dpi compared to 0 dpi (Fig 3C). There was a trend for the maximum continuous length to be greater at 1 dpi following neutrophil depletion, but the effect was not significant (Fig 3C, P = 0.18). Nevertheless, neutrophil-depleted mice exhibited greater capillary perfusion as visualized by FITC-dextran circulating in the bloodstream (Fig 3D).

**Figure 3.**
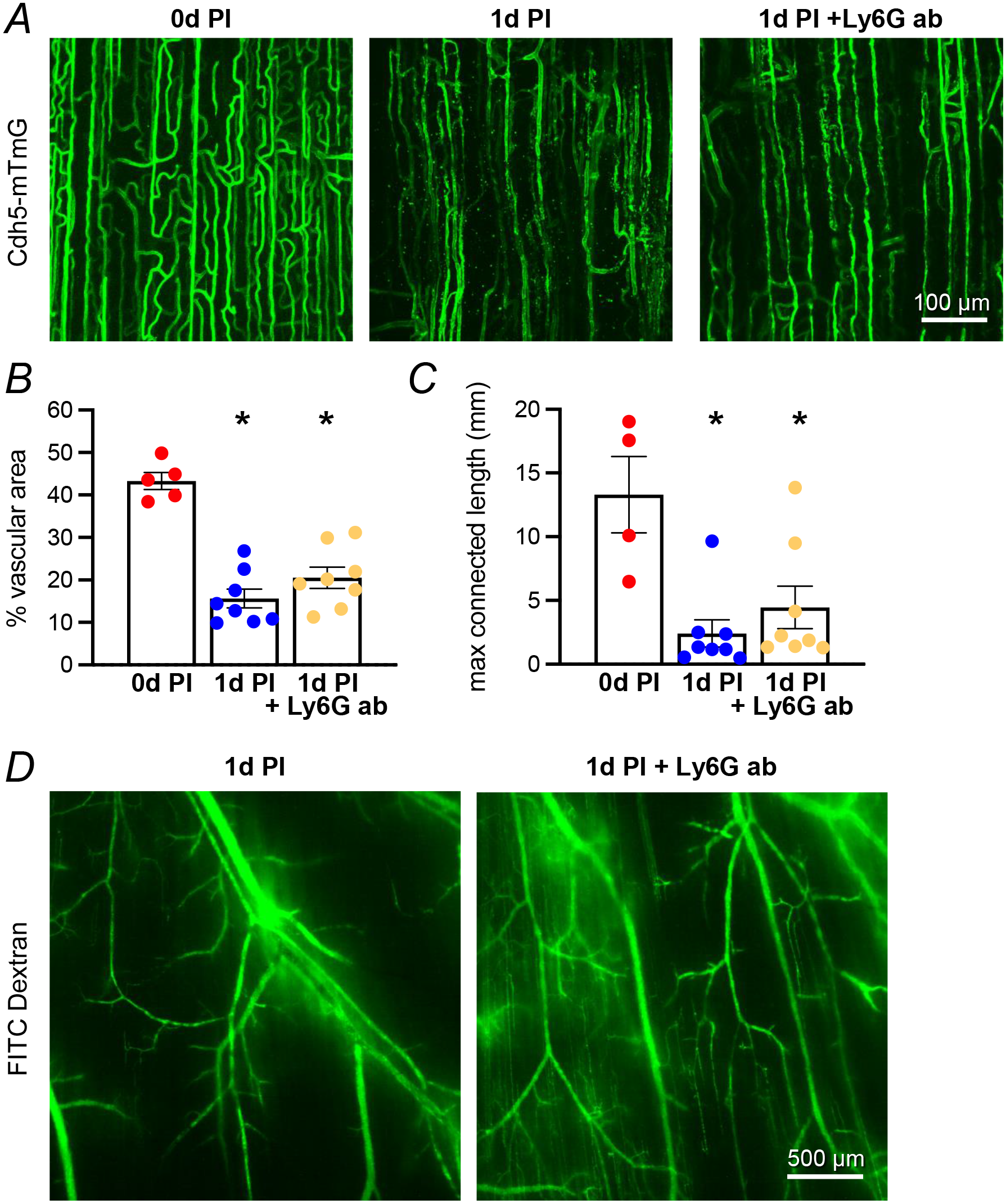
Skeletal muscle injury with BaCl_2_ disrupts capillary networks. **A)** Representative maximum projection z-stacks of GM from 4-5 Cdh5-mTmG mice. Capillary ECs (green; eGFP) are well organized and align with myofibers (oriented vertically, not shown) in uninjured muscle at 0 dpi. At 1 dpi after BaCl_2_ injection, capillaries are fragmented with few intact segments. Neutrophil depletion with Ly6G antibody (+ Ly6G ab) affords partial protection against capillary damage at 1 dpi. Scale bar applies to all panels. **B)** Muscle injury with BaCl_2_ decreases the % area occupied by capillary ECs at 1 dpi. Neutrophil depletion did not prevent capillary fragmentation at 1 dpi. *1 dpi (n = 8) and 1 dpi + Ly6G antibody (n = 8), P <0.0001 vs. 0 dpi (n = 5). **C)** Injury by BaCl_2_ decreases the maximum continuous length of capillary networks, indicating fragmentation. *1 dpi (n = 8), *1 dpi + Ly6G ab (n = 8), *P <0.01 vs. 0 dpi (n = 4). **D)** Representative images from 4 mice show intravascular FITC-dextran (green, 70 kDa) labeling of perfused residual microvascular networks with some dye leakage into surrounding tissue. At 1 dpi, more capillaries remain perfused by FITC-dextran following neutrophil depletion using Ly6G ab. Scale bar applies to both panels. Summary data in B and C are means ± SE.

### Regenerating capillary networks are dilated and disorganized

By 2-3 dpi, endothelial sprouts emerge from surviving capillary segments at multiple initiation points. The ensuing angiogenesis reestablished perfused networks at 5 dpi (Fig 4A) as reported (9). After a ∼65% reduction at 1 dpi (Fig 3B), the % vascular area recovered to uninjured levels at 5 dpi (Fig 4B) and did not change thereafter. However, capillary networks appeared disorganized and dilated during their regeneration, with diameter increasing from 4.7 ± 0.1 µm at 0 dpi to 6.9 ± 0.2 µm at 5 dpi and 6.1 ± 0.6 µm at 10 dpi (Fig 4C). By 21 dpi, capillary diameter (5.7 ± 0.3 µm) was no longer significantly different from uninjured controls.

**Figure 4.**
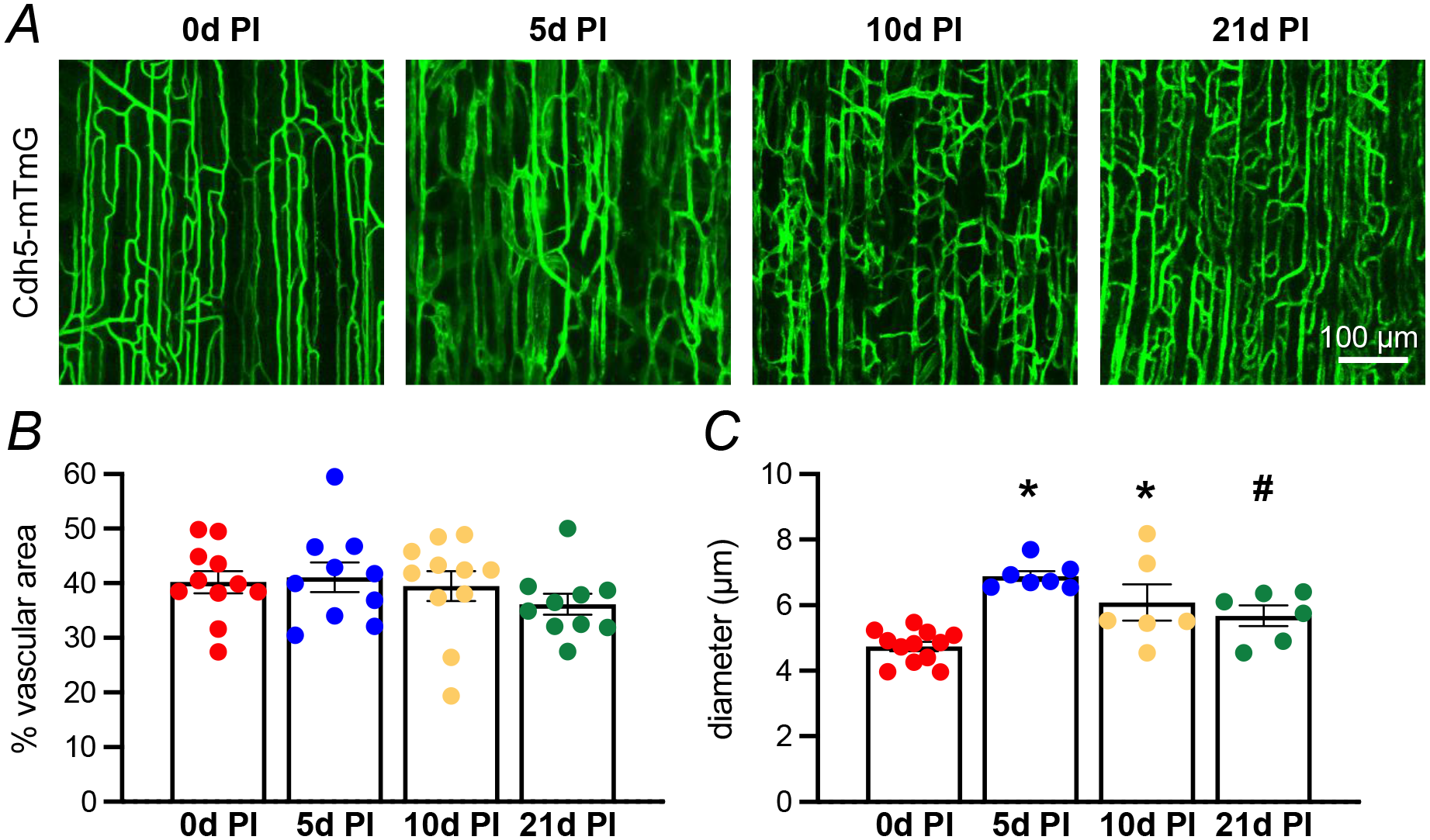
Regenerated capillaries are initially dilated when reperfused, but microvascular density is similar throughout regeneration. **A)** Maximum projections of confocal z-stacks acquired from GM of Cdh5-mTmG mice before injury and during regeneration (ECs = green; eGFP). Representative images from 4-5 mice. Scale bar applies to all panels. **B)** Following capillary fragmentation at 1 dpi (see Figure 3A), the % vascular area has returned to control levels by 5 dpi and is not different throughout regeneration (n = 10-11 images from 5-6 mice per timepoint). **C)** Capillaries are dilated at 5 and 10 dpi but return to uninjured diameter by 21 dpi (n = 6-12 images per timepoint from 4 mice) *P <0.01 vs. 0 dpi; #P <0.05 vs. 5 dpi. Summary data are means

To quantitatively examine morphological changes in regenerating capillary networks, we first evaluated the number of branch points. In uninjured muscle (0 dpi), there was 26 ± 2 branch points/mm of network length and this value did not change throughout regeneration (Fig 5A). However, capillary regeneration was nonuniform. Poorly vascularized regions (not shown) were located adjacent to regions of high angiogenic activity which often had elongating sprouts that connected with neighboring capillaries to form a dense microvascular mesh (Fig 5B) that resembled a developmental vascular plexus (22). Trifurcations not found in uninjured muscle were also present within regenerating networks at 10 dpi, albeit with low incidence (Fig 5B).

**Figure 5.**
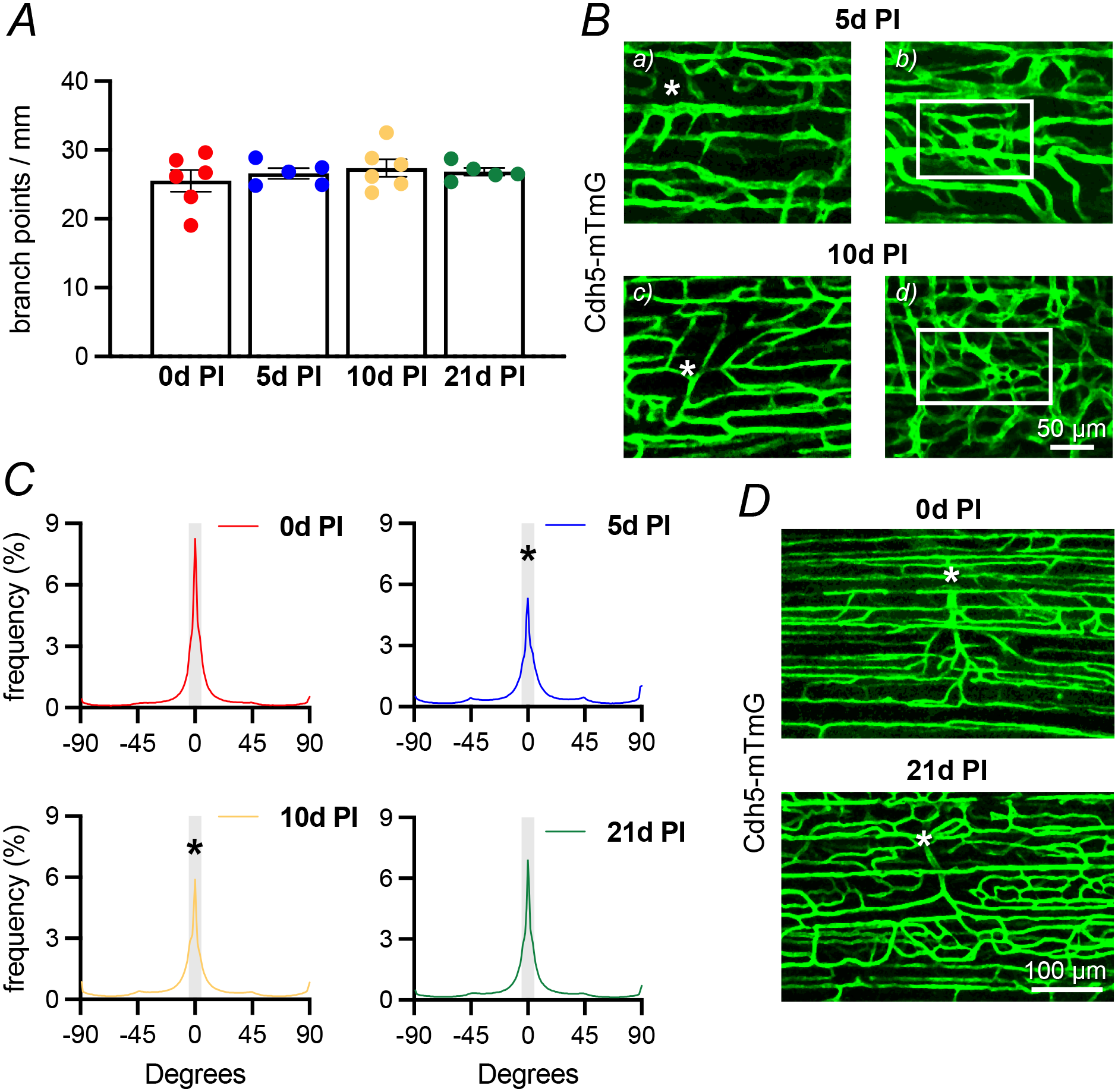
Regenerated capillary networks are disorganized. **A)** Branch point frequency (# / total capillary vascular length, mm) is not different throughout regeneration (n = 5-6 images from 4-5 mice). Summary data are means ± SE. **B)** Confocal z-stacks of GM from Cdh5-mTmG mice highlighting unique capillary structures during regeneration. At 5 dpi injury, *a)* sprouts elongate towards adjacent capillaries creating *b)* a dense mesh (within box). By 10 dpi, *c)* trifurcations (*) are present, which are not found in uninjured muscle. In addition, *d)* multiple anastomoses create plexus-like structures (box) similar to those occurring during development (ECs = green; eGFP). Scale bar applies to all panels. **C)** Regenerated capillaries are less well aligned with myofibers (oriented vertically at 0°) at 5d and 10 dpi compared to uninjured controls (0 dpi). By 21 dpi, capillaries have remodeled such that their orientation is no longer different from 0 dpi (n = 5-6 per time point from 4-5 mice, *P<0.01 vs. 0 dpi). Grey bar: ± 1 standard deviation of Gaussian fit at 0 dpi for capillaries oriented parallel to myofibers (−5 to +5°) superimposed for each timepoint for reference. **D)** A microvascular unit (MVU) constitutes a terminal arteriole (*) and the capillaries it supplies in both directions. MVUs are readily identified at 0 dpi but cannot be resolved before 21 dpi because of disoriented segments (e.g, panels in B).

Nascent capillaries grew in multiple directions such that fewer capillaries aligned with regenerating myofibers, as documented by an orientation analysis. To calculate the % of capillary segments parallel to myofibers, the distribution of their angles was plotted relative to myofibers, which were oriented vertically at 0° for reference and horizontal defined as −90° and +90°. Capillary segments in uninjured GM oriented within 1 standard deviation of vertical were considered parallel to myofibers (−5 to +5°). This interval was used as a reference for 5, 10 and 21 dpi in Figure 5. In uninjured muscle (0 dpi), 47 ± 3% of capillaries were parallel to myofibers (Fig 5C). However, at 5 dpi, only 34 ± 2% of regenerated capillaries were parallel with myofibers; this structural disorientation persisted through 10 dpi. However, by 21 dpi the % of capillaries orientated parallel to myofibers was no longer different from 0 dpi. Capillary network disorientation was manifested by the loss of identifiable microvascular units (MVUs), defined as a group of capillaries supplied by a common terminal arteriole (23). Restoration of organized MVUs characteristic of uninjured skeletal muscle was not apparent until 21 dpi (Fig 5D), which corresponds with maturation of regenerating myofibers in the mouse GM following BaCl_2_ injury (9).

### Terminal arterioles remodel during muscle regeneration

Finding minimal cellular damage in arterioles (Fig 2) but extensive remodeling of capillaries and MVUs (Fig 5), we questioned whether the architecture of arteriolar networks underwent remodeling during regeneration. At criterion timepoints, the GM vasculature of male C57BL/6J mice was manually traced from the inferior gluteal artery (feed artery) to terminal arterioles (Fig 6). Resulting traces included intra-network anastomoses and collateral connections to arterioles originating from the superior gluteal artery.

**Figure 6.**
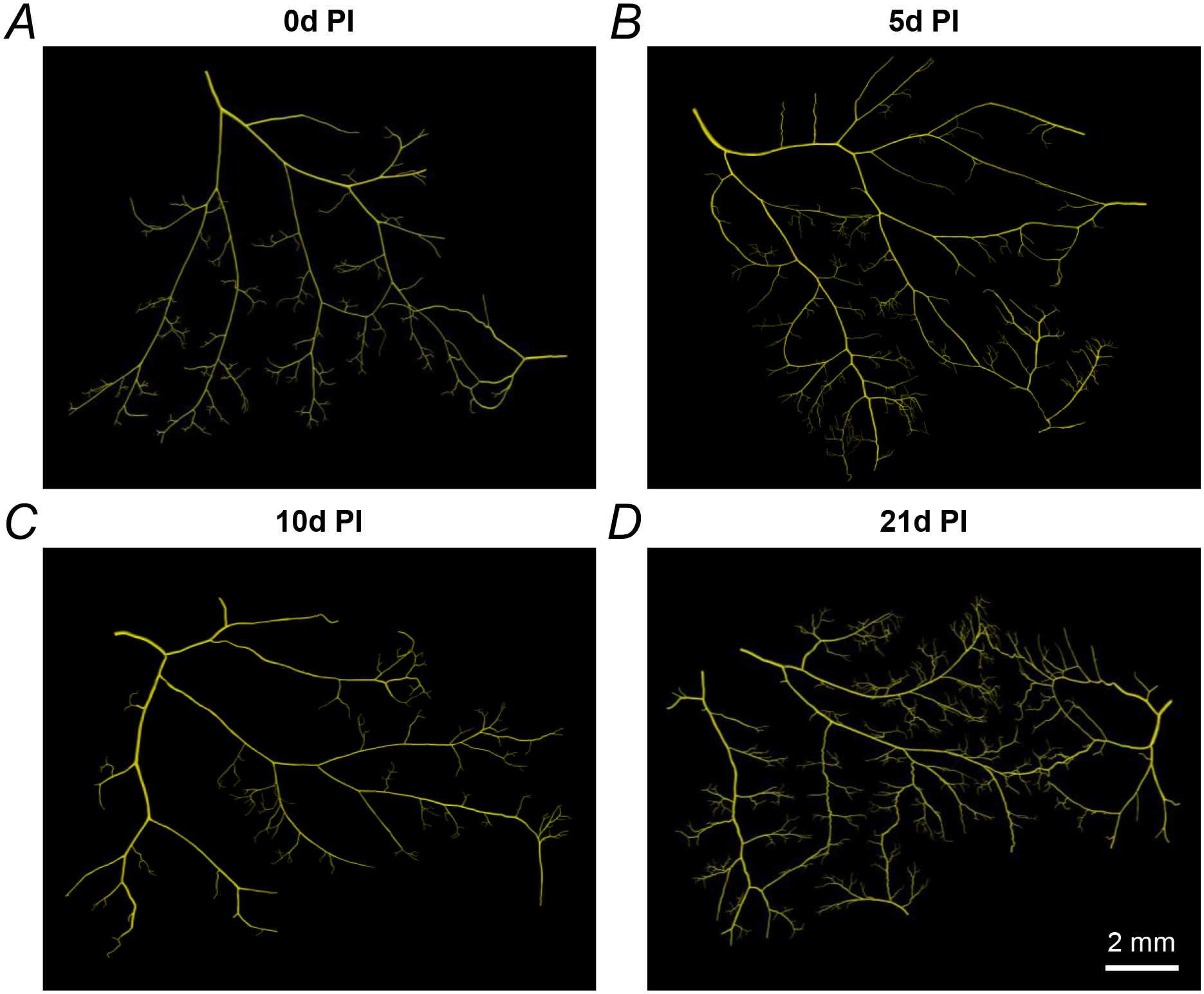
Representative maps of resistance networks from feed artery to terminal arterioles in **A)** uninjured GM (0 dpi) and at **B)** 5, **C)** 10, and **D)** 21 dpi. Vascular networks from 5-6 GM were reconstructed in Vesselucida software. Scale bar applies to all panels.

Uninjured muscle contained 452 ± 78 microvessel segments (Fig 7A). During regeneration, the total # of segments was not different compared to 0 dpi; the total vascular length was also not different at the timepoints studied. However, when categorized by microvessel diameter, the # of arterioles 5-10 µm in diameter (i.e., terminal arterioles) increased at 5 dpi compared to 0 dpi and remained elevated through 21 dpi (Fig 7B). The corresponding cumulative vascular length of the smallest arterioles was also increased at 5 dpi vs. 0 dpi indicating proliferation of terminal arterioles (Fig 7C). In addition, more anastomoses were present in microvascular networks at 5 dpi than other timepoints (Fig 7D), reflecting the rapid angiogenesis during the early stages of regeneration.

**Figure 7.**
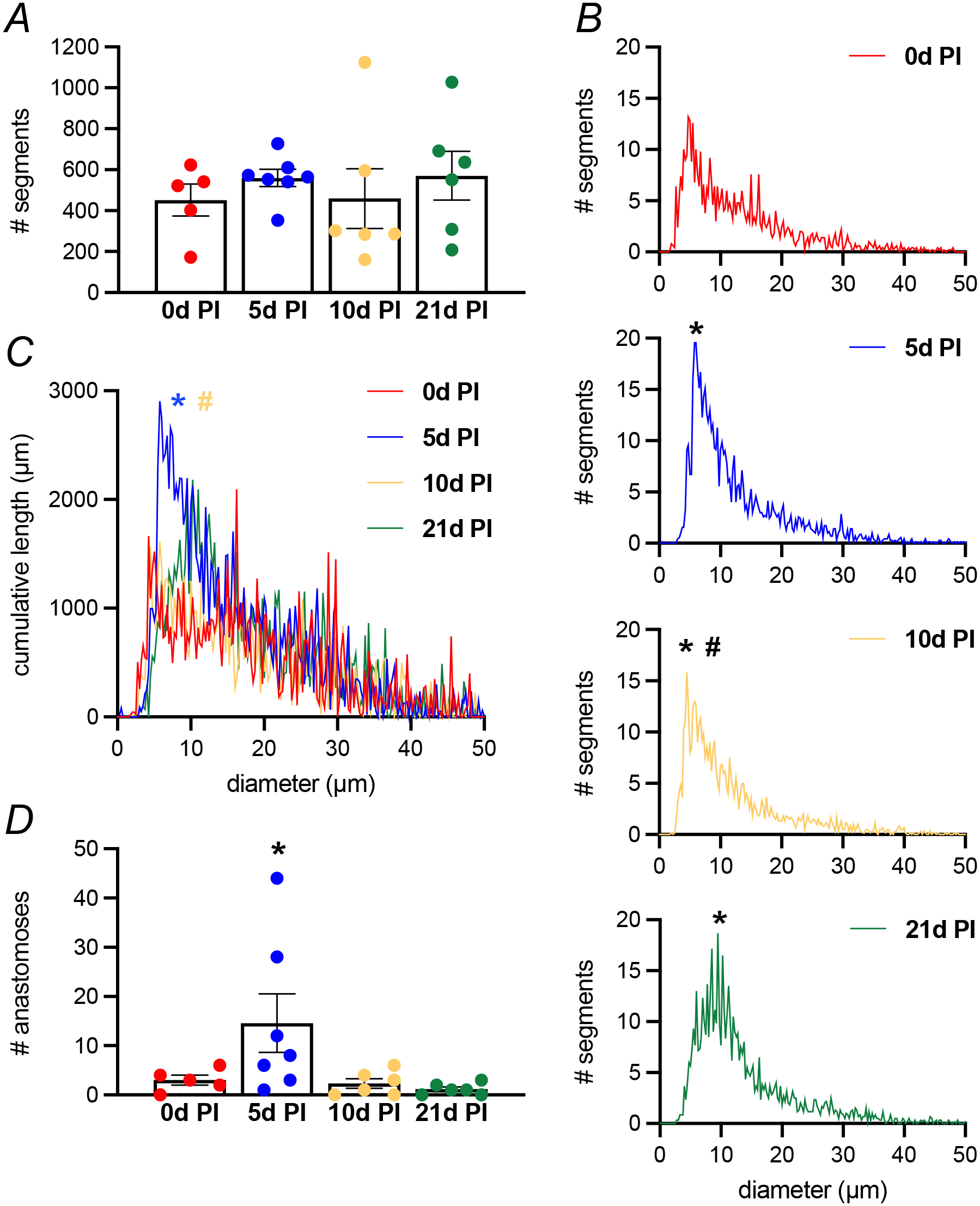
Terminal arterioles in resistance networks proliferate after muscle injury. **A)** The total number of segments in resistance networks is not significantly different between uninjured GM and during regeneration (n = 5-7 GM per timepoint, as depicted in Fig 6). **B)** Segments 5-10 µm diameter proliferate at 5 dpi, which persists through 21 dpi (n = 5-7 GM per timepoint). Error bars removed for clarity; *P<0.05 vs. 0 dpi, ^#^P<0.05 vs. 5 dpi. **C)** The cumulative length of terminal arterioles in GM at 5 dpi is increased compared to uninjured GM (n = 5-7 GM per timepoint_. Error bars not shown for clarity. *P<0.05 vs. 0 dpi, ^#^P<0.05 vs. 5 dpi. **D)** Anastomoses are more prevalent in arteriolar networks at 5 dpi compared to 0 dpi. By 10 dpi, the # anastomoses returned to uninjured levels (*P<0.05 vs. 0 dpi, n = 5-6 GM per timepoint). Summary data are means ± SE.

## Discussion

Apart from myofibers, microvascular cells represent the largest cell population in healthy adult skeletal muscle (24). This relationship underscores the reliance of skeletal muscle on a robust microvascular supply to support the metabolic demands of myofibers. Because myofibers depend upon their microvascular supply for survival, understanding how microvascular cells respond after skeletal muscle injury and during regeneration is integral to the development of therapies targeted to promote muscle regeneration.

### Microvascular cell damage after muscle injury

We showed that BaCl_2_ induces depolarization of SMCs and ECs (Fig 1A & B), consistent with its action as a broad-spectrum K^+^ channel inhibitor (25, 26). However, the consequences of such an effect differ between microvascular cells and myofibers. In myofibers, BaCl_2_-induced depolarization is accompanied by a rise in [Ca^2+^]_i_ resulting in proteolysis, membrane disruption, and cell death within 1 h (8). Elevation of [Ca^2+^]_i_ can increase mitochondrial Ca^2+^ content to trigger cytochrome C release and intrinsic apoptosis (27). For SMCs exposed to the same conditions, membrane depolarization also increased [Ca^2+^]_i_, which led to vasoconstriction (Fig 1D), as seen in rabbit aorta (28). However, since cell death did not occur after the same duration of exposure (or even when extended to 3 h), the present findings indicate that either longer exposure to BaCl_2_ is necessary (which is unlikely given the equal probability for BaCl_2_ diffusion into both cell types) or that the rise in [Ca^2+^]_i_ was not sufficient to induce SMC death. Finding negligible disruption of arteriolar or venular SMCs at 1 dpi (Fig 2) when myofibers have degenerated supports the latter explanation. We suggest that varying amounts of mitochondria, expression of different L-type Ca^2+^ channel isoforms, the ability for the sarcoplasmic/endoplasmic reticulum to sequester Ca^2+^ without dysfunction, or expression of pro-apoptotic proteins may explain the difference in cell death between myofibers and microvascular SMCs exposed to BaCl_2_ (29, 30).

When recording from ECs in endothelial tubes, depolarization occurred without an increase in [Ca^2+^]_i_ (Fig 1B &C), which is consistent with their lack of voltage operated Ca^2+^ channels in the plasma membrane (13). Furthermore, no cell death was evident in endothelial tubes exposed to BaCl_2_ (Fig 1E). That ECs comprising capillaries are damaged following BaCl_2_ injection *in vivo* [Fig 3A, (7, 8)], but ECs directly exposed to BaCl_2_ *in vitro* are not (Fig 1E), suggests that ECs in arteriolar and venular networks may be protected by SMCs *in vivo* and that the otherwise exposed capillary ECs are injured secondary to myofiber degeneration from BaCl_2_ exposure. Upon exposure to BaCl_2_, resting force of the mouse EDL muscle increases ∼40% over 15 min, which then returns to baseline as proteolysis disrupts the integrity of contractile proteins (8). Given that evidence of capillary damage is not apparent during this time, the present data imply that skeletal muscle tissue degeneration induces capillary fragmentation hours after injury, rather than as a consequence of persistent myofiber contraction.

Within 1-2 h of damage by BaCl_2_ or physical trauma, skeletal muscle is invaded by neutrophils that generate pro-inflammatory cytokines, chemokines, and reactive oxygen species which exacerbate tissue damage (20). While neutrophil depletion prior to BaCl_2_ injection did not prevent capillary damage at 1 dpi (Fig 3A & B), more capillaries remained perfused when compared to mice in which neutrophils were not depleted (Fig 3D). This outcome suggests that while neutrophil activity contributes to capillary damage, additional mechanisms lead to EC death when skeletal muscle is injured. Candidates include toxic substances released from degenerating myofibers, reactive oxygen species, calpain-dependent protein degradation, recruitment of proinflammatory cells including monocytes and macrophages (20, 31–34).

### Restoration of capillary networks during muscle regeneration

Previous studies investigating capillarity during regeneration evaluated muscle cross sections (7, 35) and therefore cannot address the dynamics of microvascular network organization. Cross sections are inherently biased towards vessels oriented parallel to myofibers (36), which compromises the resolution of transverse microvessels and anastomoses. Moreover, studies that have examined the intact microcirculation during skeletal muscle regeneration have focused on hindlimb muscles such as the tibialis anterior (7) and EDL (10), which restricts observations to the superficial portion of the muscle due to the thickness of the tissue. While capillaries, terminal arterioles, and collecting venules can be observed, proximal resistance networks reside deeper in the muscle. To overcome these limitations, we evaluated microvascular regeneration in the GM, a thin, flat skeletal muscle well suited to high resolution imaging and analysis of entire microvascular networks (9).

Triggered by the release of such growth factors as VEGF, FGF, and IGF-1 from the hypoxic milieu (37), nascent capillaries sprout and elongate from surviving microvessel fragments at 2-3 dpi (7, 8). These regions of angiogenic activity within the tissue highlight the selective roles of individual ECs during angiogenesis. As shown during development, endothelial tip and stalk cells are selected after stimulation by VEGF and Notch signaling (38). Tip cells guide capillary sprouts as they sense attractive and repulsive signals in the microenvironment. Stalk cells trail tip cells and proliferate to elongate nascent capillaries such that by 5 dpi, capillary networks have reformed (Fig 4A) and perfusion is restored (9). While others have reported no change or even a reduction in the diameter of capillaries during the early stages of muscle regeneration (7), we observed a ∼30% increase in capillary diameter at 5 and 10 dpi (Fig 4B). Vasodilation is one of the earliest steps in both physiological and pathological angiogenesis (39). That resistance arterioles are also dilated at this time (9) suggests that regenerating skeletal muscle requires elevated blood flow afforded by a dilated microvasculature providing ample nutrients for cellular regeneration. Capillary dilation may reflect aberrant coverage by mural cells (pericytes) or dysregulated inter- and intracellular signaling (40–42).

Following injury of the GM, adjacent to sites of angiogenesis and regeneration are regions with extensive damage that are poorly perfused or avascular at 5 dpi, implying that microvascular regeneration after skeletal muscle injury is asynchronous between neighboring regions of tissue. Heterogeneity amongst ECs exists between vascular beds, microvessels, and even along the length of a single microvessel in healthy skeletal muscle (43, 44). Corresponding differences in gene expression and nuances in EC function may explain the asynchronous regenerative response of the microvasculature. In response to tissue hypoxia, angiogenesis occurs primarily in areas containing glycolytic (type II) myofibers (45, 46), suggesting that diffusion distances for O_2_ and metabolites affect the angiogenic response. That the GM is a muscle comprised of mixed fiber type (47) is consistent with the observed heterogeneity in angiogenesis during regeneration of the GM.

In addition to the density of capillary networks, optimal distribution of capillaries is crucial for adequate muscle oxygenation. Heterogenous capillary spacing negatively impacts muscle oxygenation due to local increases in diffusion distances (48, 49). Microvascular structures resembling a developmental vascular plexus were present throughout the GM at 5 and 10 dpi (Fig 5B), with disorganized capillary networks containing abnormal branching (e.g., trifurcations) and more segments that were not aligned with regenerating myofibers (Fig 5C). Similar abnormal microvascular morphogenesis has been described after grafting the hamster tibialis anterior muscle (50) or ischemic injury to mouse EDL muscle (10).

A microvascular unit constitutes a terminal arteriole and the group of capillaries it supplies (3). Following injury of the GM, the organization of capillaries into discernable MVUs apparent in uninjured muscle was not identifiable until 21 dpi (Fig 5D), the timepoint which coincides with the recovery of blood flow control in the resistance vasculature of the GM following BaCl_2_ injury (9). In uninjured healthy skeletal muscle, terminal arterioles are oriented obliquely to myofibers, whereas capillaries primarily run parallel with myofibers. When a terminal arteriole constricts or dilates, flow through all capillaries it supplies diminishes or increases, respectively. We anticipated that nascent microvessels would repopulate residual basement membranes (51–54) following microvascular damage from skeletal muscle injury (7, 55). In contrast to the organization of MVUs in healthy muscle, regenerated microvascular networks were less aligned with myofibers, contained irregular structures, and required 21 dpi to approximate normal structure. This delay in reorganization implies that nascent capillaries did not regrow along basement membranes of old microvessels. We suggest that these structural abnormalities in capillary networks contribute to nonuniform perfusion during the early stages of myofiber regeneration. The persistence of arteriolar dilation during this time maintains capillary perfusion, which may help compensate for limitations imposed by aberrant capillary network organization.

### Remodeling of arteriolar networks during regeneration

Muscle injury by BaCl_2_ did not induce vascular cell death or structurally damage the resistance (or adjacent venular) network upstream from terminal arterioles (Fig 2). Evaluating network architecture during regeneration revealed an increase in the number of the smallest (i.e., terminal) arterioles 5-10 µm in diameter that persisted through 21 dpi (Fig 7B &C). A similar increase in small arterioles was observed after ischemic injury in the mouse spinotrapezius muscle after arterial ligation (56). We also observed an increase in the number of anastomoses in arteriolar networks at 5 dpi (Fig 7D). These direct connections ensure redundancy in blood flow paths to limit underperfusion in addition to promoting the maximum delivery and removal of blood to regenerating tissue before local blood flow control recovers. While the origin of new arterioles in microvascular networks during regeneration remains incompletely understood, nascent capillaries may become “arterialized” by recruiting mural cells (57). Together, the data suggest that the repair, recovery, and maintenance of new myofibers requires a robust supply of oxygen and nutrients from the regenerating microcirculation preceding the recovery of blood flow regulation.

### Conclusion

Skeletal muscle is highly vascularized. Capillaries are located parallel to and in close association with myofibers, providing oxygen and nutrients while removing metabolic byproducts. The present study shows that capillaries, but not SMCs or ECs of larger microvessels, are disrupted by the microenvironment created by degenerating myofibers. When perfusion is restored, nascent capillary networks are dilated, disorganized, and associated with more terminal arterioles. These early morphological adaptations may compensate for lack of blood flow regulation during myofiber regeneration (9). Reorganization of capillaries and terminal arterioles into MVUs at 21 dpi coincides with restoration of the number myofibers, their cross-sectional area, and the recovery of blood flow regulation in resistance networks. Understanding how microvascular structure and function are restored following skeletal muscle injury provides new insight for developing therapeutic interventions for the treatment of acute muscle trauma in the adult.

## Materials and Methods

### Ethical approval

All procedures were approved by the Animal Care and Use Committee at the University of Missouri (protocol #10050) and were performed in accordance with the National Research Council’s Guide for the Care and Use of Laboratory Animals and the animal ethics checklist of this journal.

### Animal care and use

Male C57Bl/6J mice were purchased from Jackson Laboratory (Bar Harbor, ME, USA) at ∼14 weeks of age and acclimated at the University of Missouri animal care facilities at least 1 week prior to study. Male Cdh5-mTmG mice [cross of VE-cadherin-CreERT2 mice (21) (gifted from Dr. Luisa Iruela-Arispe) and Rosa26-mTmG mice (#007676, Jackson Laboratory); both on C57BL/6 background] were bred and housed in animal care facilities of the University of Missouri. Mice were studied at ∼4 months of age. Cre recombination for membrane-bound eGFP expression in ECs was induced through intraperitoneal injection of 100 µg tamoxifen (1 mg/100 µL in peanut oil; #T5648, Sigma-Aldrich; St. Louis, MO, USA) on 3 consecutive days with at least 1 week allowed after the first injection prior to study. All mice were maintained under a 12:12 h light/dark cycle at 22-24°C with fresh food and water *ad libitum*. To control for an order effect, criterion time points and treatment status were randomized.

### Preparation of isolated microvessels and endothelial tubes

On the morning of an experiment, a male C57Bl/6J mouse was anesthetized [ketamine (100 mg/kg) + xylazine (10 mg/kg) in sterile saline; intraperitoneal injection], abdominal fur was shaved, and a midline incision through the skin was made from the sternum to the pubis. The abdominal muscles were exposed, removed bilaterally, and placed in a dissection chamber containing chilled, nominally Ca^2+^ free physiological salt solution (PSS, pH 7.4) containing (in mM): 140 NaCl (Fisher Scientific; Pittsburgh, PA, USA), 5 KCl (Fisher), 1 MgCl_2_ (Sigma), 10 HEPES (Sigma), and 10 glucose (Fisher); standard PSS also contained 2 mM CaCl_2_ (Fisher). Muscles were pinned as a flat sheet onto transparent silicone rubber (Sylgard 184; Dow Corning; Midland, MI, USA). While viewing through a stereomicroscope, an unbranched segment of the superior epigastric artery [SEA; length, ∼2 mm; diameter, ∼150 µm; comprised of a single smooth muscle cell (SMC) layer surrounding the endothelial cell (EC) monolayer] was dissected from the surrounding tissue. Following isolation, an SEA was transferred to a tissue chamber (#RC27-N; Warner Instruments; Hamden, CT) for cannulation. The tissue chamber was secured in a platform with micromanipulators (MT-XYZ; Siskiyou Corp; Grants Pass, OR, USA) positioned at each end that held heat-polished cannulation micropipettes (external diameter, ∼100 µm). The SEA was cannulated at each end and secured with suture. The vessel preparation was transferred to the stage of a Nikon E600FN microscope (Tokyo, Japan) mounted on a vibration isolation table (TMC Vibration Control; Peabody, MA, USA). The vessel was pressurized to 100 cm H_2_O (∼75 mmHg), maintained at 37°C, superfused at 3 mL/min with standard PSS, and allowed to equilibrate for 15 min before experimentation.

To isolate intact endothelial tubes, an SEA segment (length, ∼1 mm, diameter, ∼60 μm) was placed into a round bottom test tube containing 0.62 mg/mL papain (#P4762, Sigma), 1 mg/mL dithioerythritol (#D8255, Sigma), and 1.5 mg/mL collagenase (#C8051, Sigma) in PSS and incubated for 30 min at 34°C (11). The vessel segment was transferred to the tissue chamber and gently triturated to remove the SMCs by aspirating and ejecting the segment through borosilicate glass capillary tubes that were heat polished at one end (tip internal diameter, ∼80 μm). Following dissociation of SMCs (confirmed by visual inspection at 200X magnification), the endothelial tube was secured against the bottom of the chamber with blunt fire-polished micropipettes held in the micromanipulators. The preparation was secured on an inverted microscope (#TS100, Nikon) mounted on a vibration isolation table (TMC Vibration Control; Peabody, MA, USA). The endothelial tube was maintained at 33°C, superfused at 3 mL/min with standard PSS, and equilibrated for 15 min before experimentation.

### Intracellular recording

Membrane potential (V_m_) of SMCs (intact pressurized SEA) or ECs (freshly isolated endothelial tube) was recorded with an Axoclamp amplifier (2B; Molecular Devices; Sunnyvale, CA, USA) using micropipettes pulled (P-97; Sutter) from glass capillary tubes (#GC100F-10; Warner; Hamden, CT, USA) and backfilled with 2 M KCl (tip resistance, ∼150 MΩ). A Ag/AgCl pellet was placed in effluent standard PSS for the reference electrode. The output of the amplifier was connected to a data acquisition system (Digidata 1322A; Molecular Devices) and an audible baseline monitor (ABM-3; World Precision Instruments; Sarasota, FL, USA). Data were recorded at 1000 Hz using Axoscope 10.1 software (Molecular Devices) on a personal computer. Successful impalements were indicated by sharp negative deflection of V_m_, stable V_m_ > 1 min, and prompt return to 0 mV upon withdrawal of the electrode. Once a cell was impaled, V_m_ was recorded for at least 5 min to establish a stable baseline. The superfusion solution was then changed to PSS containing 1.2% BaCl_2_ until the V_m_ response had stabilized (∼10 min). Each experiment represents paired data under resting baseline conditions and when stabilized during with BaCl_2_ treatment for an intact vessel (SMCs) or endothelial tube (ECs) from a separate mouse.

### Calcium photometry

A cannulated, pressurized SEA was secured in a tissue chamber and placed on an inverted microscope. The vessel was superfused (3 mL/min) at 37°C for 20 min with standard PSS, then incubated in a static bath containing fura 2-AM dye (#F14185, Fisher). The dye was dissolved in DMSO, diluted to 1 µM in standard PSS, and added to the tissue chamber for 40 min. Superfusion with standard PSS was then resumed for 20 min to wash out excess dye. Fura 2 fluorescence was used to evaluate [Ca^2+^]_i_ by alternatively exciting the preparation at 340 nm and 380 nm while recording emissions at 510 nm through a 20X objective [Nikon Fluor20, numerical aperture (NA) = 0.45] using IonWizard 6.3 software (IonOptix, Milford, MA). After fluorescence and vessel diameter were recorded under baseline conditions, 1.2% BaCl_2_ was added to the superfusion solution. Intracellular Ca^2+^ signals and vessel diameter were measured over 30 min of BaCl_2_ exposure. Under these conditions (dye loaded from the bath), the [Ca^2+^]_i_ signal primarily originates from SMCs (16).

To measure [Ca^2+^]_i_ in ECs, an endothelial tube was incubated for 30 min with fura 2-AM dye in a static bath and then washed for 20 min with standard PSS. To maintain their integrity, these preparations were studied at 33°C (58). After baseline fluorescence was recorded, Ca^2+^ signals (F_340_/F_380_) were measured during 30 min of exposure to 1.2% BaCl_2_ in standard PSS; responses typically stabilized within ∼10 min.

Each experiment represents paired data under resting baseline conditions and during stabilization of the response to BaCl_2_ treatment for an intact vessel or endothelial tube from a separate mouse.

### Cell death

Following equilibration in standard PSS, the superfusion solution was changed to PSS containing 1.2% BaCl_2_. After preparations were exposed to BaCl_2_ for 1 h [which kills >90% myofibers, (8)], superfusion with standard PSS was restored. The membrane permeant nuclear dye Hoechst 33342 (1 µM; #H1399, Fisher) was used to identify nuclei of all cells and propidium iodide (2 µM, #P4170, Sigma) to identify nuclei in dead and dying cells (14, 15). Following BaCl_2_ exposure, respective nuclear dyes (in standard PSS) were perfused through the lumen of a cannulated SEA (0.1 mL/min) or superfused over the surface of an endothelial tube for 10 min followed by 10 min wash in standard PSS.

To evaluate cell death (14), fluorescent images of nuclear staining with Hoechst 33342 and propidium iodide were acquired with appropriate filters using a 40X water immersion objective (NA = 0.8) coupled to a DS-Qi2 camera with Elements software (version 4.51) on an E800 microscope (Nikon). Z-stacks were acquired from the top half of the vessel segment and analyzed using ImageJ software. Stained nuclei were counted manually within a defined region of interest (150 x 500 µm in intact microvessels, 50 x 300 µm in endothelial tubes); nuclei of ECs are oval shaped and oriented parallel to the vessel axis while SMC nuclei are thin and oriented perpendicular to the vessel axis (14). For each cell type, cell death is expressed as a percentage as follows: (# propidium iodide^+^ nuclei / # Hoechst 33342^+^ nuclei) x 100.

### Muscle injury by local injection of BaCl_2_

A mouse was anesthetized with ketamine + xylazine and rested on an aluminum warming plate to maintain body temperature. The skin over the injury site was shaved and sterilized by wiping with Betadine Solution (Purdue Products LP; Stamford, CT, USA) followed by 70% alcohol. An incision (∼5 mm) was made through the skin to expose the gluteus maximus muscle (GM) near the lumbar fascia. Using a Hamilton syringe and 32 gauge needle (Reno, NV, USA), 75 µL of 1.2% BaCl_2_ in water was injected under the GM to injure the muscle (9). The incision was closed with surgical glue or two discontinuous sutures. The mouse was placed on a heated platform, monitored until consciousness and ambulation were restored, then returned to its original cage. Mice routinely recovered normal activity and behavior within 24 h and were studied up to 21 dpi with uninjured mice (0 dpi) serving as controls. The 21 dpi time point coincides with recovery of vasomotor control in arteriolar networks and restoration of myofiber number during regeneration of the mouse GM following BaCl_2_ injury (9).

### Neutrophil depletion

Inflammation is integral to muscle injury (20). In some experiments, neutrophils were depleted prior to BaCl_2_ injection using a neutralizing Ly6G antibody. Mice were injected intraperitoneally with either 500 μg of anti-Ly6G 1A8 antibody [#BE0075, BioxCell; Lebanon, NH, USA; (59)] or vehicle (sterile saline) on −2, −1, and 0 dpi. Neutrophil depletion was confirmed by a differential blood count of a sample collected by cardiac puncture in mice anesthetized with ketamine + xylazine. A drop of whole blood was spread on a glass slide and stained with a Wright Giemsa stain. Leukocytes were counted in sets of 100 cells, differentiating between lymphocytes, neutrophils, and monocytes. Two sets of 100 counts were averaged per slide. Blood samples were obtained from mice without muscle injury (0 dpi) and at 1 dpi and compared to mice injected with the vehicle at the same time points.

### Dissection of gluteus maximus muscle

A mouse was anesthetized with ketamine + xylazine as above. Supplemental doses (∼20% of initial dose) were given (intraperitoneal injection) throughout the experimental protocol (typically lasting 2-3 h) to maintain a stable plane of anesthesia (checked every 15 minutes by lack of withdrawal to tail or toe pinch). Hair was removed by shaving and the mouse was placed on a warming plate. As previously described (60), skin and connective tissue overlying the GM were removed using microdissection to expose the GM. Once exposed, the GM was continuously superfused with a bicarbonate-buffered physiological salt solution (bb-PSS; pH 7.4, 34-35°C) containing (in mM) 131.9 NaCl_2_ (Fisher), 4.7 KCl (Fisher), 2 CaCl_2_ (Sigma), 1.17 MgSO_4_ (Sigma), and 18 NaHCO_3_ (Sigma) equilibrated with 5% CO_2_/95% N_2_. While viewing through a stereomicroscope, the GM was cut along its origin from the lumbar fascia, sacrum, and iliac crest to reflect the muscle away from the body and reveal its vascular supply on the ventral side. The muscle was then either removed for immunostaining or prepared for intravital microscopy as described below.

### Whole mount immunostaining for confocal imaging

The GM from C57BL/6J mice was excised, placed in ice-cold phosphate buffered saline (PBS, pH 7.4; #P3813, Sigma) and placed onto transparent rubber coated 12 well plate. The muscle was spread to approximate its dimensions *in situ* with the ventral surface facing up and secured at the edges with insect pins. Excessive connective tissue and fat were removed using microdissection. The GM was fixed overnight at 4°C in 4% paraformaldehyde in PBS. After washing in PBS (3 x 5 min), the muscle was immersed in blocking buffer containing 0.5% Triton x-100 (#T8787, Sigma), 2% bovine serum albumin (#BP671; Fisher), and 4% normal goat serum (#50197Z, Sigma) in PBS. To identify SMCs, the GM was stained with monoclonal rabbit anti-Myh11 [1:500; #ab124679; Abcam, Cambridge, UK; (61)] in blocking buffer overnight at 4°C, then washed with blocking buffer (3 x 5 min), incubated with goat anti-rabbit Alexa 633 (1:200; #A21071, Fisher) in blocking buffer for 2 h at room temperature, and washed in PBS (3 x 10 min).

An immunostained GM was placed in a custom imaging chamber with the ventral surface facing the objective to optimize resolution of the microvasculature. A small volume of PBS (<10 µL) was added to the chamber and the GM was flattened by placing a glass block (2 cm x 2.5 cm x 1 cm; mass, 7.8 g) on the dorsal surface. Images of the microvasculature were acquired with a 10X objective (NA = 0.4; image size, 1162.5 µm x 1162.5 µm) or 25X water immersion objective (NA = 0.95; image size, 465 µm x 465 µm) on a laser scanning confocal microscope (TCS SP8, Leica Microsystems; Buffalo Grove, IL) using Leica LASX software. Images were digitally rotated such that myofibers were aligned vertically for analysis of capillary orientation. Maximum projection z-stacks (z-step, 2 µm; ∼150 µm thick) were used to resolve SMC coverage and capillary network morphology (n = 4-5 mice, 5-12 images across mice).

### Analysis of capillary diameter and network morphology

Male Cdh5-mTmG mice expressing green fluorescent protein (eGFP) in ECs were used to analyze capillary diameters and network morphology during regeneration. The GM was excised, cleaned, and imaged as described above, but without prior fixation. To measure the average diameter of capillaries from confocal z-stacks (image size, 465 µm x 465 µm), a calibrated 5×5 grid was centered over the image, creating 25 equal ROIs (93 µm x 93 µm). The diameter of 5 capillaries were measured manually from 10 of 25 ROIs selected at random; these 50 measurements were averaged for the image with 2-3 images analyzed per muscle from 4-6 mice. To analyze capillary network morphology using ImageJ, a threshold was applied to maximum projections of z-stacks to generate a binary image. The % vascular area (area fraction) was calculated by determining the area occupied by eGFP fluorescence within the entire image (465 µm x 465 µm).

To assess capillary network morphology from a larger field of view (1162.5 µm x 1162.5 µm), the binary image was skeletonized and analyzed with the Analyze Skeleton plugin (ImageJ). Outputs included: # segments, length of segments, # connected segments, and # branch points. Segments included in this analysis contained at least 3 continuous pixels, which corresponded to ∼2.5 µm.

To calculate the continuous capillary length as a measure of fragmentation at 1 dpi, the length of connected segments was summed for each network, defined as a group of connected capillaries, and the maximum continuous capillary length determined for a single image (465 µm x 465 µm); the mean ± SE of all images (n = 4-8 images from 4 mice) are reported. Branch point frequency was calculated as the number of branch points per total microvascular length (mm) for a given image. The OrientationJ plugin generated distribution histograms for capillary orientation analysis with −90° and +90° on the horizontal axis. All measurements in ImageJ were validated using reference networks created with known area, length, number of branch points, and orientation.

### Intravital microscopy

The GM of anesthetized C57BL/6J mice was dissected as described above, then spread onto a transparent rubber pedestal and pinned at its edges to approximate *in situ* dimensions (9, 60). The preparation was transferred to the stage of a Nikon E600FN microscope and equilibrated for 30 min while continuously superfused with bb-PSS at 3 mL/min maintained at 34-35°C (pH, 7.4). To assess vascular perfusion and permeability during maximum capillary damage (1 dpi), a fluorescein isothiocyanate (FITC) conjugated dextran (70 kDa, to approximate the mass of albumin) was injected into the retro-orbital sinus to access the systemic circulation and allowed to circulate for ∼10 min. The preparation was illuminated by a mercury lamp for fluorescence imaging using appropriate filters. Images were acquired through a 4X objective (Nikon Fluor4; NA = 0.1; image size, 2.7 mm x 3.4 mm) coupled to a low light CMOS FP-Lucy camera (Stanford Photonics; Palo Alto, CA, USA) and displayed on a digital monitor. Time lapse images were recorded at 40 frames/s using Piper Control software (Stanford Photonics). At the end of the experiment, the mouse was given an overdose of anesthetic and killed by cervical dislocation.

### Mapping arteriolar networks

In anesthetized mice, wheat germ agglutinin conjugated to Alexa Fluor 647 (WGA- 647, 1 mg/mL, 200 µL; #W32466, Fisher) was injected into the retroorbital sinus to access the systemic circulation and label the endothelial glycocalyx. The WGA-647 distributed throughout the vascular compartment for 10 min. Thereafter, the GM was dissected, cleaned, and secured with pins as above. The GM was fixed overnight at 4°C (4% paraformaldehyde in PBS) avoiding direct light. Tissues were washed in PBS, cleared in 100% glycerol overnight at 4°C, then rinsed in PBS, mounted onto a slide, and coverslipped.

Whole mount GM preparations were viewed using a Nikon Eclipse 600 microscope with a Nikon Plan Fluor 10X objective (NA 0.3) coupled to a CMOS camera (Orca Flash 4.0; Hamamatsu) and personal computer. The XYZ translational stage (Ludl Electronic, Hawthorne, NY, USA) was controlled by stepper motors coupled to an integrated joystick for constant repositioning of the tissue during imaging. The tracing icon was adjusted by the operator to match vessel diameter; spatial resolution was < 2 µm. This strategy is similar to that used in aged mice (60). A grid slide (MBF Bioscience; Williston, VT) was used for parcentric and parfocal calibration when linking the hardware with the software. A stage micrometer (100 × 0.01 = 1 mm; Graticules Ltd, Tonbridge Kent, UK) validated the calibration.

### 3D reconstruction and analysis of resistance networks

Vesselucida Microscope Edition software (MBF Bioscience) was used to trace entire resistance networks in the GM, from the inferior gluteal (feed) artery to terminal arterioles. Some tracings contained collateral branches to the resistance network fed by the superior gluteal artery. Occasionally, in thicker regions of the GM, reduced visibility resulted in discontinuous or incomplete networks, which were not used in the present analyses. Furthermore, for criterion data, networks containing fewer than 100 vessel segments in the completed tracing were excluded from analyses. Vesselucida Explorer software (MBF Bioscience) was used to calculate # segments, diameter, length, and # of anastomoses in each tracing. To generate summary histograms across GM preparations during regeneration, segments were binned into diameter increments of 1 µm; for cumulative length, those within each bin were summed. Anastomoses were defined as the shortest length to return to a given branch point.

### Statistics

Statistical analyses were performed using Prism 9 software (GraphPad Software Inc., La Jolla, CA, USA). For *in vitro* experiments of isolated vessel preparations, V_m_, [Ca^2+^]_i_, and diameter were analyzed with a paired two-tailed Student’s t-test. For capillary network morphology, variables evaluated during regeneration were analyzed by 1-way ANOVA and Tukey’s multiple comparisons post-hoc test. Frequency distributions for dimensions of resistance networks were analyzed with a Kolmogorov-Smirnov test. P < 0.05 was considered statistically significant. Values for n given in figure captions refer to the number of vessels or images evaluated from 4-6 mice.

## Acknowledgements

The authors thank Dr. Robert Arpke for performing tamoxifen injections for Cdh5-mTmG mice. Dr. Erika Boerman provided use of her confocal microscope for imaging capillary network morphology. Dr. Susan Tappan and Timothy Tetreault at MBF Bioscience provided expert technical assistance with applying Vesselucida software to our analyses of arteriolar networks.

## Additional Information

### Funding

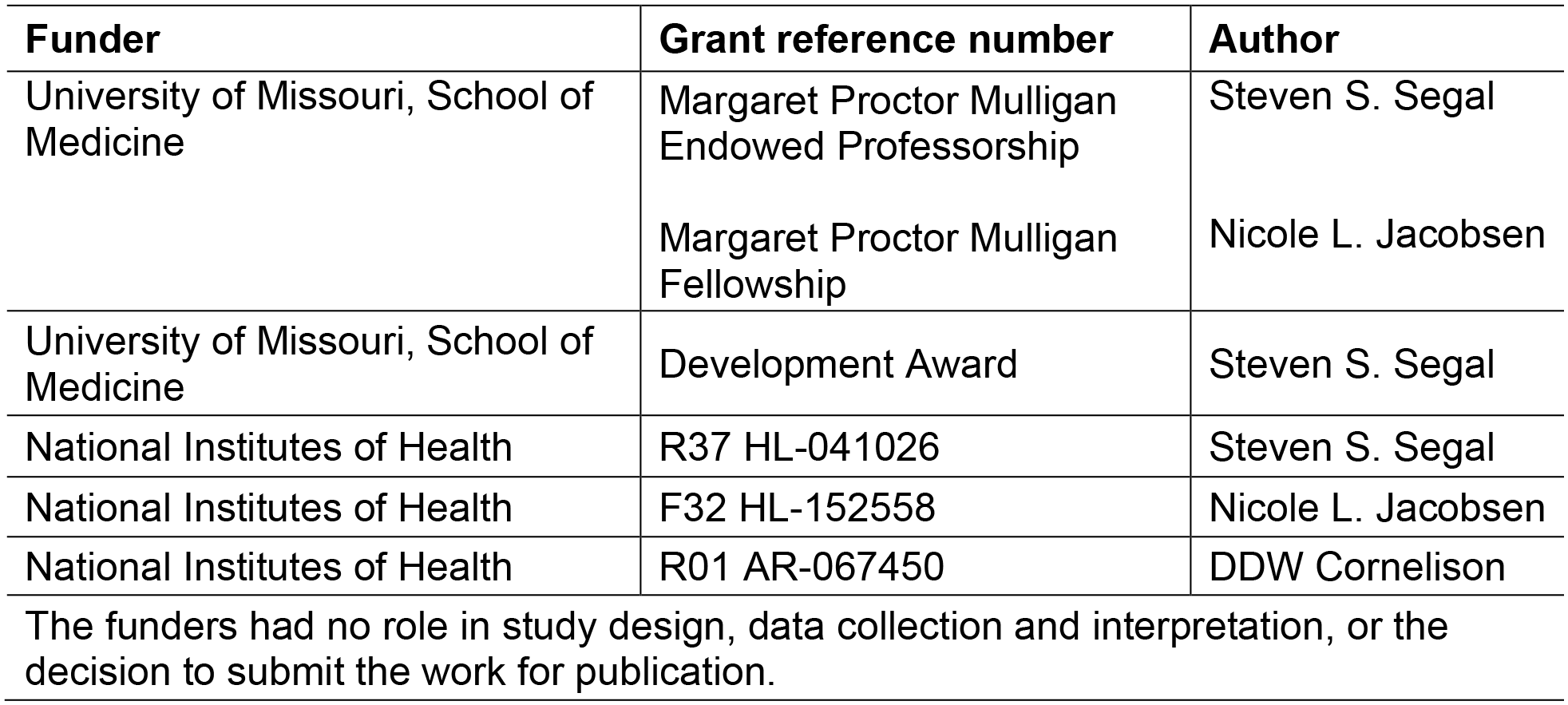

| **Funder** | **Grant reference number** | **Author** |
| --- | --- | --- |
| University of Missouri, School of Medicine | Margaret Proctor Mulligan Endowed Professorship | Steven S. Segal |
|  | Margaret Proctor Mulligan Fellowship | Nicole L. Jacobsen |
| University of Missouri, School of Medicine | Development Award | Steven S. Segal |
| National Institutes of Health | R37 HL-041026 | Steven S. Segal |
| National Institutes of Health | F32 HL-152558 | Nicole L. Jacobsen |
| National Institutes of Health | R01 AR-067450 | DDW Cornelison |
| The funders had no role in study design, data collection and interpretation, or the decision to submit the work for publication. |  |  |

### Author Contributions

NL Jacobsen: Conceptualization, Data Curation, Formal Analysis, Validation, Investigation, Methodology, Funding Acquisition, Writing-original draft, Writing-review and editing; CN Norton: Data Curation, Formal Analysis, Validation, Investigation, Methodology, Writing-review and editing; RL Shaw: Data Curation, Investigation, Methodology, Writing-review and editing; DDW Cornelison: Conceptualization, Supervision, Funding Acquisition, Resources, Writing-review and editing; SS Segal: Conceptualization, Data Curation, Supervision, Methodology, Funding Acquisition, Resources, Writing-review and editing, Project Administration.

